# An extensive archaeological dental calculus dataset spanning 5000 years for ancient human oral microbiome research

**DOI:** 10.1101/2024.09.17.613443

**Authors:** Francesca J. Standeven, Gwyn Dahlquist-Axe, Jessica Hendy, Sarah Fiddyment, Malin Holst, Krista McGrath, Matthew Collins, Amy Mundorff, Anita Radini, Josef Wagner, Conor J. Meehan, Andrew Tedder, Camilla F. Speller

**Affiliations:** School of Chemistry and Biosciences, University of Bradford; BioArCh, Department of Archaeology, University of York, York, UK; York Osteoarchaeology Ltd.; Department of Prehistory and Institute of Environmental Science and Technology (ICTA-UAB), Universitat Autònomade Barcelona, Bellaterra, Spain; Section for Evolutionary Genomics, the GLOBE Institute, University of Copenhagen, København, Denmark; Department of Archaeology, University of Cambridge, Cambridge; Department of Anthropology, College of Arts and Sciences, University of Tennessee, Knoxville, TN, USA; UCD School of Archaeology, College of Social Sciences and Law, University College Dublin, Dublin, Republic of Ireland; Enteric Diseases, Murdoch Children’s Research Institute, Parkville, Australia; Department of Paediatrics, The University of Melbourne, Parkville, Australia; Department of Biosciences, Nottingham Trent University; Department of Anthropology, University of British Columbia

## Abstract

Archaeological dental calculus can provide detailed insights into the ancient human oral microbiome. We offer a multi-period, multi-site, ancient shotgun metagenomic dataset consisting of 174 samples obtained primarily from archaeological dental calculus derived from various skeletal collections in the United Kingdom. This article describes all the materials used including the skeletons’ historical period and burial location, biological sex, and age determination, data accessibility, and additional details associated with environmental and laboratory controls. In addition, this article describes the laboratory and bioinformatic methods associated with the dataset development and discusses the technical validity of the data following quality assessments, damage evaluations, and decontamination procedures. Our approach to collecting, making accessible, and evaluating bioarchaeological metadata in advance of metagenomic analysis aims to further enable the exploration of archaeological science topics such as diet, disease, and antimicrobial resistance (AMR).

## Background

Ancient oral microbiomes have provided a wealth of information about past lives by analysing dental calculus (mineralised plaque) —a common archaeological material recovered from skeletons spanning most pre-historic and historical periods (Forshaw 2022). Dental calculus is a mineralized biofilm, formed initially by an acquired enamel pellicle (AEP) on the tooth surface, followed by bacterial colonisers, and a series of microbial succession. As dental calculus mineralises, it entombs and preserves biomolecules associated with oral microbiota (Adler et al. 2013; Huynh et al. 2016; Jersie-Christensen et al. 2018; Granehäll et al. 2021), environmental and dietary micro debris (Warinner et al. 2014b; Cristiani et al. 2016; Weyrich et al. 2017; Sawafuji et al. 2020), and the human host —although present in consistent but low quantities (Mann et al. 2018). Previous metagenomic analyses of dental calculus have demonstrated a range of archaeological applications, including: microbiota changes with ancient dietary shifts (Adler et al. 2013); sugar intake in archaeological populations (Chidimuro et al. 2022); early domesticated food consumption (Cristiani et al. 2016); historical human migratory events (Eisenhofer et al. 2019); the presence of various pathogens (Huynh et al. 2016; Fotakis et al. 2020; Jacobson et al. 2020; Neukamm et al. 2020); the identification of dietary sources indicated through ancient proteins (Hendy et al. 2018); the tracing of agricultural transitions (Ottoni et al. 2021); and evidence of ancient dairy consumption (Warinner et al. 2014a).

This dataset is intended for research capitalising on the abundant microbial aDNA encapsulated in ancient dental calculus (Metcalf et al. 2014; Brealey et al. 2020) to gain potential insights into millennia of oral microbial evolution. Our sizable dataset allows for the comparison of oral microbiomes across multiple historic time periods in England (Bronze/Iron Age, Roman, Anglo-Saxon/Viking, Medieval, Industrial, Post-Industrial, modern) spanning 2500 BC to 1900 AD (and 20^th^ century). This time span encompasses significant historical events such as the agricultural revolution, industrialisation, and medical discoveries, which permit the analysis of changes in dietary, demographic, environmental, and socioeconomic aspects over time. For example, recent studies have identified the Industrial Revolution as a period of shifting community function (Gancz et al. 2023) and diversity (Skelly 2019); yet datasets including ancient oral metagenomes dating to this period are sparse. Therefore, this dataset focuses on expanding the availability of metagenomes dating to the period of cultural and environmental transition during the Industrial Revolution in England, as well as samples from earlier and later periods that contextualise the Industrial Era. The inclusion of populations from both urban centres and rural communities enables research into the impact of geographic differences on ancient oral microbiomes, as well as other environmental factors associated with industrialisation.

Our published dataset facilitates the study of oral microbiomes of burial populations, and/or single individuals, which can contribute to debates in funerary archaeology, social construction, religious history, and health disparities between different social classes. For example, previous research has demonstrated important population-level data that can be retrieved from sites like Spitalfields, London (Adams and Reeve 1987; Molleson et al. 1993; Nitsch et al. 2011; Humphrey et al. 2012), Iron Age and Romano-British settlements (Oliver 1992; Pearce 1999; Watts 2001; Redfern et al. 2015), cemeteries such as the *Cillinis* that entomb groups of unbaptized children (Finlay 2000), and plague pits (Papagrigorakis et al. 2007; Harbeck et al. 2013; Wagner et al. 2014; Barbieri et al. 2017; Spyrou et al. 2018). Nevertheless, every ancient individual is a unique entity of time deserving of their own exploration, and previous research has also demonstrated novel insights that can be obtained from the analysis of solitary individuals, for example: Otzi the Iceman (Acs et al. 2005; Keller et al. 2012; Seiler et al. 2013; Lugli et al. 2017; Larcher 2019; Pires de Souza et al. 2021); the human genome that was retrieved from a piece of prehistoric Scandinavian birch pitch (Jensen et al. 2019); and ‘bog bodies’ (Nielsen et al. 2020; van Beek et al. 2023).

Our dental calculus dataset also makes a thorough analysis of the interactions between various microbial groups possible. Diseases in the oral cavity are caused by the dysbiosis of a diverse array of microbiome members, rather than by a single pathogen (Almeida et al. 2020). For example, the correlation of carbohydrate diets with *Streptococcus mutans* formation and its visible effects on ancient teeth is intriguing (Adler et al. 2017); but holobiont units are considerably more complex than that. While *S. mutans* are metabolising sugars, producing acids, and decaying teeth, they form plaque biofilms that bind other hazardous bacteria together and spread to other areas of the oral cavity and body (Zayed et al. 2021). *S. mutans* is just an example, but our oral microbiome data offers the chance to investigate the relationships of other possible multiple microbial communities that have the potential to reveal links between past diseases and dietary patterns, in addition to current health challenges.

In addition to ancient dental calculus, our dataset also includes metagenomic data retrieved from associated archaeological tooth roots and skeletal elements which act as controls to ensure robust and reliable data can be obtained. The inclusion of environmental controls, such as soil from the burial site or bone or tooth root from a skeleton from the same archaeological site, allows for ancient and modern environmental contaminants to be identified and removed (Kazarina et al. 2021). There is currently a scarcity of literature exploring the specific impact of environmental controls on ancient oral metagenomes, as well as very few examples of their use in ancient dental calculus studies. Preliminary studies have indicated that ancient environmental contaminants can be a particular issue in metagenomic studies as their DNA damage patterns can mirror those of the targeted oral microbial DNA (Kazarina et al. 2021). Failure to thoroughly decontaminate ancient oral metagenomes is known to artificially inflate alpha diversities (Farrer et al. 2021) and therefore the inclusion of environmental control is an important consideration. Further research into the impact of environmental controls is necessary.

## Methodology

### Summary of all materials

All our samples have been archived in the European Nucleotide Archive (ENA 2023) as fastq files in projects PRJEB1716, PRJEB12831, and PRJEB75938 (see **Table S1**). They encompass 174 samples (348 fastq files using paired end reads) from modern (*n=10*) and archaeological dental calculus (*n=133*) and tooth (*n=2*) samples, environmental controls (bone; *n=7*), and laboratory controls (extraction and library blanks; *n=22*). The ancient samples were excavated from multiple archaeological sites across the UK spanning 5000 years, including the Bronze Age (ca. 2300 BC–700 BC), Iron Age (ca. 700 BC–43 AD), Roman Period (ca. 43 AD–410 AD), Anglo-Saxon Period (ca. 410 AD–1066 AD), Viking Age (ca. 793 AD–1066 AD), Medieval Period (ca. 1066–1485), Industrial (ca. 1750–1850), Post-industrial (ca. 1837–1901). Samples from modern human remains are derived from the University of Tennessee donated skeletal collection within the Forensic Anthropology Centre Anthropology Research Center (20^th^ – 21^st^ century). As an ectopic growth, dental calculus is not considered a human tissue, and analysis is not subject to the legislation of the Human Tissue Act. All samples have received ethical approval for destructive analysis and data publication from relevant repositories. All archaeological sites and time periods are displayed in **Figure 1**. **Table S1** is a summary table with all the necessary information on the individuals in our dataset including: sample code and type; skeletal ID, age, and biological sex; ENA project and number; type of tooth and weight of calculus sampled; adapter sequence; location and time period of archaeological site; and current repository and skeletal report/reference associated with individual skeletons. Proteomic analysis of a subset of these samples have also been published in Hendy et al. (2018) (see **Table S1**).

**Figure 1.**
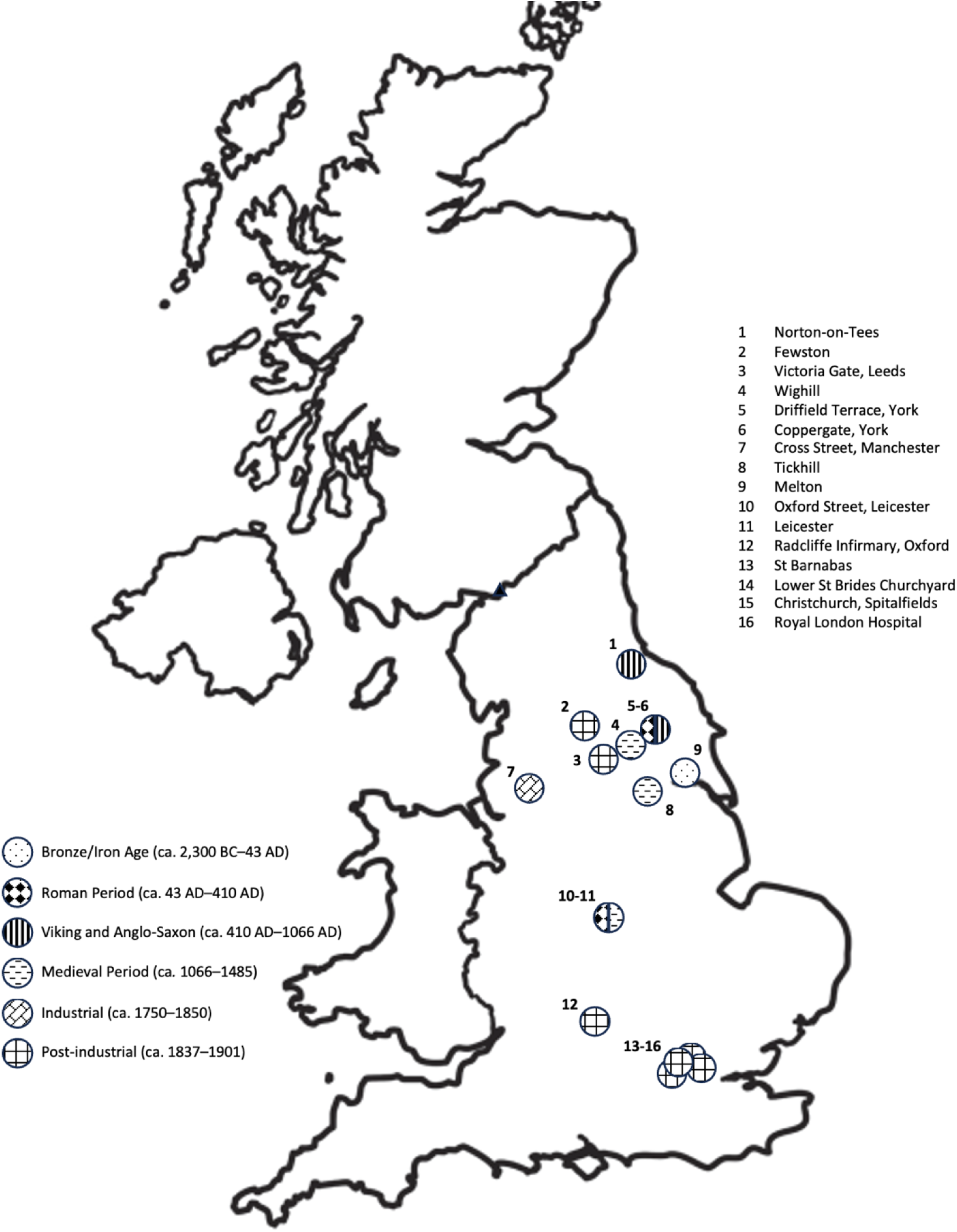
Map of dated archaeological sites across the UK from which the samples came from. More information on the exact number of samples from each site, skeletal ID, ENA accession number, and publication reference can be accessed in the supplementary data. Sample FW53C, originating from the Church of St. Michael and St. Lawrence, was not placed on the map as it could not be accurately dated. While the other samples from the site were determined to be Victorian/post-industrial, the skeletal report for FW53C did not provide a clear date. Map outline was provided by Twinkl’s free education resources (Twinkl 2023) and adapted from Hendy *et al*. 2018.

### Laboratory method

#### Sample preparation and contamination controls

Samples of dental calculus were removed from the teeth using sterilised dental picks and stored in individual 2.0 mL Eppendorf tubes. Sample preparation and DNA extractions were conducted in the BioArCh ancient DNA laboratory at the University of York, and the Ancient DNA and Proteins facility at the University of British Columbia. Both labs are dedicated to the analysis of ancient biomolecules, and the introduction of contamination into the workspace is minimised by the use of protective clothing, including Tyvek suits, gloves, masks, and hairnets. The labs are also equipped with UV filtered ventilation and positive airflow, as well as with dedicated equipment and bench UV lights; countertops and other surfaces in the lab are routinely wiped down with dilute sodium hypochlorite. All reagents and equipment in the ancient DNA laboratory are dedicated solely to the study of degraded DNA. Multiple blank DNA extractions and negative PCR controls are run alongside the ancient samples to identify potential contamination at each stage of the procedure.

#### aDNA extraction

DNA was extracted in batches, with two extraction blanks prepared alongside each batch. Samples of dental calculus and bone were UV-sterilised for 1 minute on each side. After crushing to a powder, samples were pre-digested for 5 minutes with 1 mL of 0.5M EDTA to remove possible surface contamination. This pre-digestion supernatant was removed, and a further 1.1 mL of 0.5M EDTA added and rotated at room temperature for seven days to fully demineralize. Samples were centrifuged at 13,000 RPM for 2 minutes and 1 mL of supernatant transferred into fresh tubes. To the supernatant, 20 mg/ml Proteinase K was added and rotated at 37 °C for 24 hours. For the majority of samples, DNA was extracted from the dental calculus and bone samples using a protocol based on Dabney et al. (2013) with DNA eluted in 60 µL of EB following a five-minute incubation step. For the Manchester, Cross Street samples (C028–C027), DNA extraction followed a modified silica-spin column protocol (Yang et al. 1998), with DNA concentrated in Amicon® 10K Ultra-4 Centrifugal Filter Devices (Millipore) and purified with QiaQuick MinElute kits (QIAGEN, Hilden, Germany) before being eluted in 25 µL of Qiagen EB Buffer. All DNA extracts were quantified via Qubit® 2.0 Fluorometer using a High-Sensitivity DNA Assay.

#### Metagenomic sequencing

For each DNA extract, double-stranded whole genome shotgun Illumina libraries were prepared using a protocol based on Meyer and Kircher (2010). Each dental calculus library was built using between 200-400 ng of DNA; extraction blanks were prepared with 25 µL of DNA extract; library blanks were built using 25 µL of nuclease free water. The libraries were constructed using a double-barcoding approach as described in Fortes and Paijman (2015) which serves as an additional means to filter chimeric sequences from the dataset, and thus increase the confidence in assigning the sequences to a particular library. Individual P7 indexes were ligated through an indexing PCR step using a proof-reading taq polymerase (AccuPrime Pfx Supermix) with the following cycling conditions were: 95°C for 5 minutes, and cycles of (95°C for 15s, 60°C for 30s, 68°C for 30s), and a final extension of 68°C for 5 min. Optimal cycle numbers for library indexing were determined through the use of quantitative PCR (qPCR) using Fast SYBR (Gansauge and Meyer 2013). Amplified libraries were subsequently purified using Qiagen MinElute spin columns, the size distribution of the amplified libraries was determined using an Agilent 2100 Bioanalyzer. The dental calculus libraries were pooled in equimolar concentration and subjected to paired-end sequencing on multiple HiSeq2500 lanes at the Wellcome Trust Sanger Institute (WTSI) or on a NextSeq platform, PE 150 + 150 bp Integrated Microbiome Resource (IMR) at Dalhousie (Manchester Cross Street). In accordance with WTSI protocols, human DNA sequences were removed from dental calculus datasets prior to deposition in the ENA. For samples sequenced at the WTSI (**Table S1**), the raw metagenomic reads were mapped against the human reference genome (GRCh37) using *bwa aln* with default settings; reads mapping to the human genome reads were extracted from the dataset and only unmapped reads uploaded to the ENA. Samples sequences at the IMR (Manchester Cross Street) did not have human reads filtered prior to deposition.

### Quality control

FastQC v0.11.9 (Babraham Bioinformatics 2019) was used to assess raw digital data quality. FastQC is a quality control tool for raw sequencing data that provides a modular collection of analyses used to gain insight into any flaws in the data before performing further analysis (Babraham Bioinformatics 2019). The preprocessing programmer Fastp v0.23.2 (Chen *et al*. 2018) was then utilised with default parameters. Fastp is a tool used to filter and trim poor quality reads, cut adapters, repair mismatched base pairs, and produce overall quality. It also provides results that include both pre- and post-filtering data, allowing for a direct comparison on the filtering impact (Chen *et al*. 2018).

### Decontamination

To minimise the impact of laboratory and environmental contamination, BBduk (Joint Genome Institute 2023) was used with default parameters to decontaminate samples using Kmers. This procedure involved identifying and removing homologous sequences found in extraction and library blanks, and bone blanks when they were available (see **Table 1** for which samples were paired with which controls). After decontamination, paired read counts did not match because the tool removed a different number of sequences it considered to be contaminants from forward and reverse reads, resulting in unequal counts. Therefore, the samples were subsequently processed with Trimmomatic v0.39 (Bolger et al. 2014) to re-pair reads.

**Table 1.**
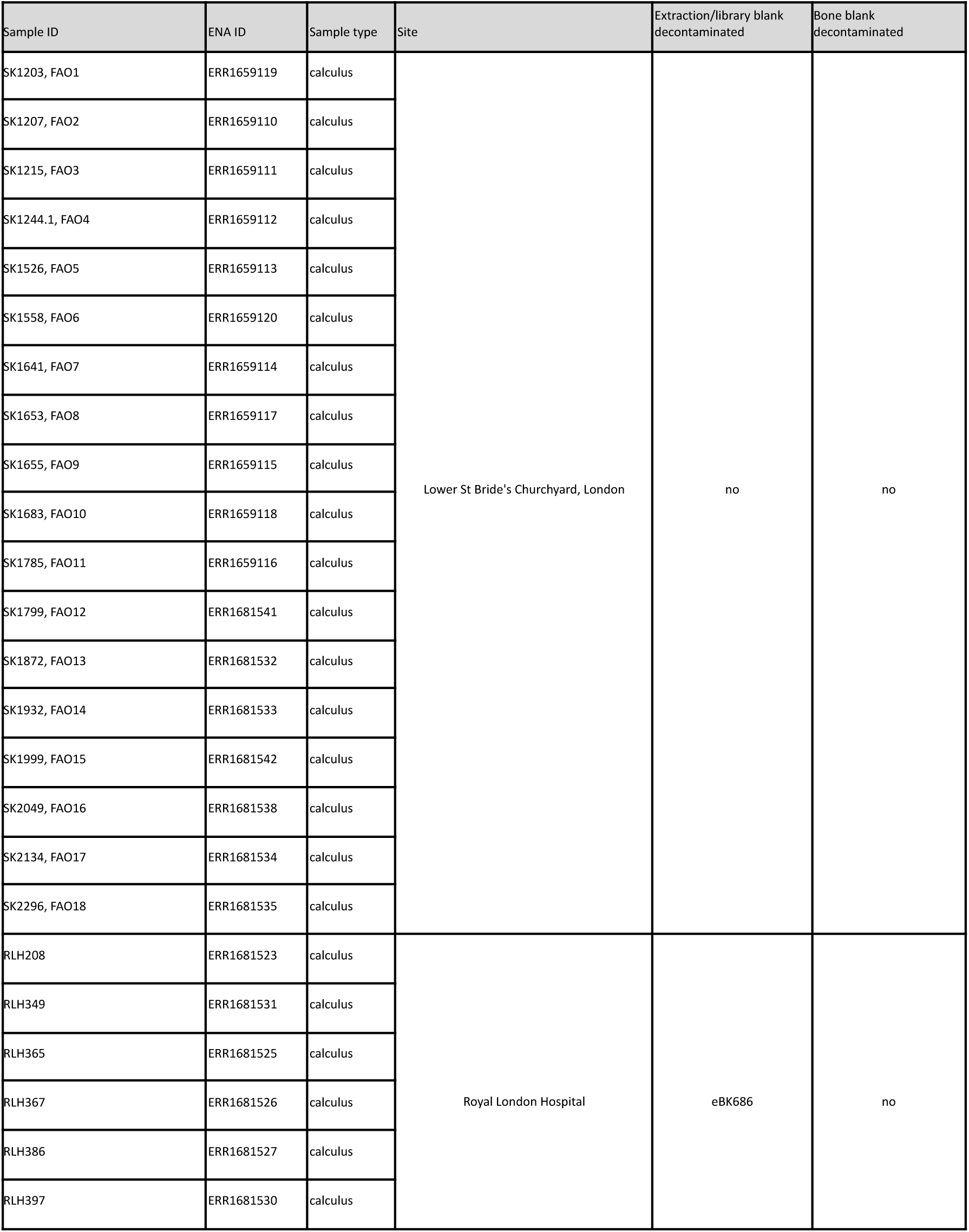
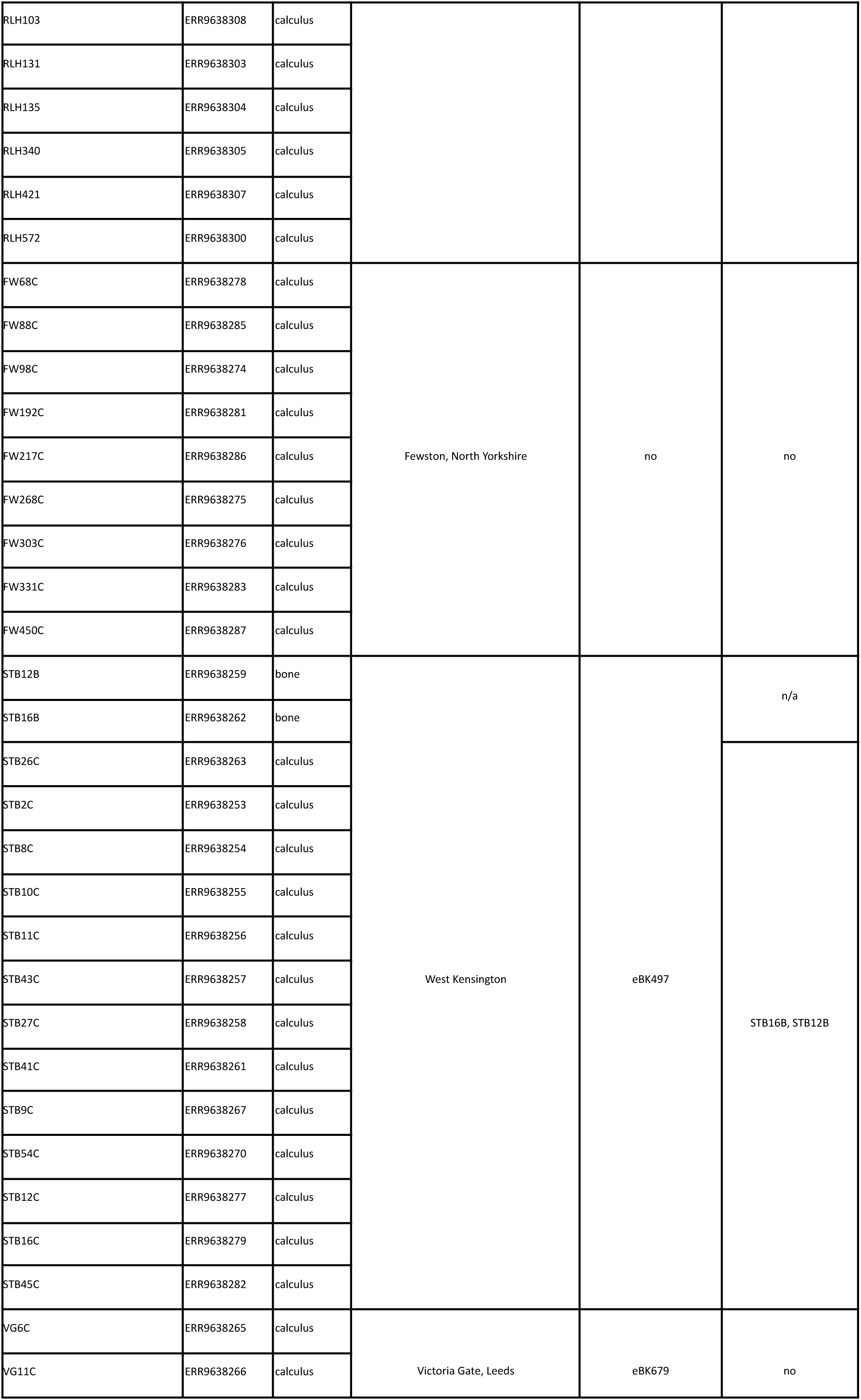
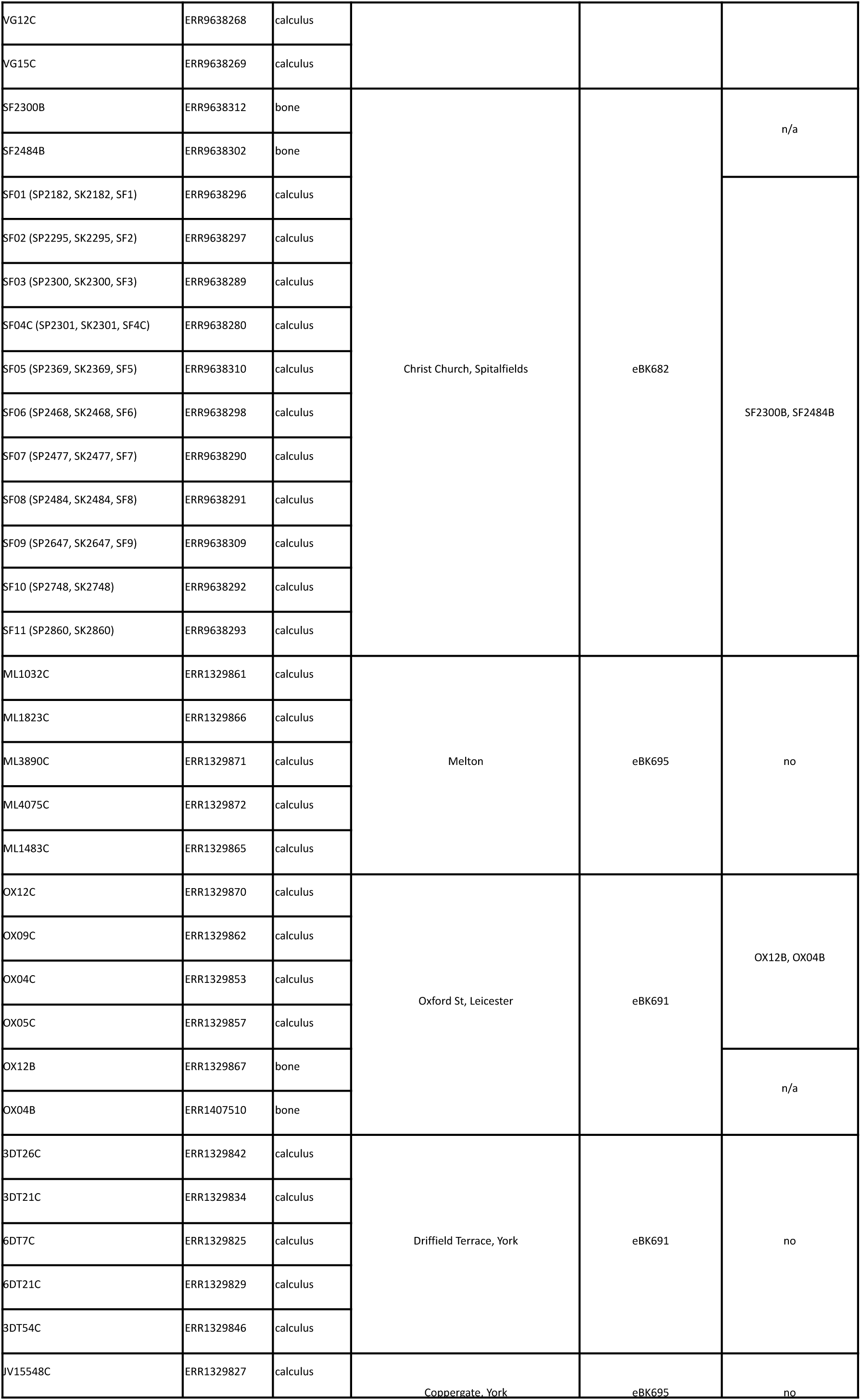
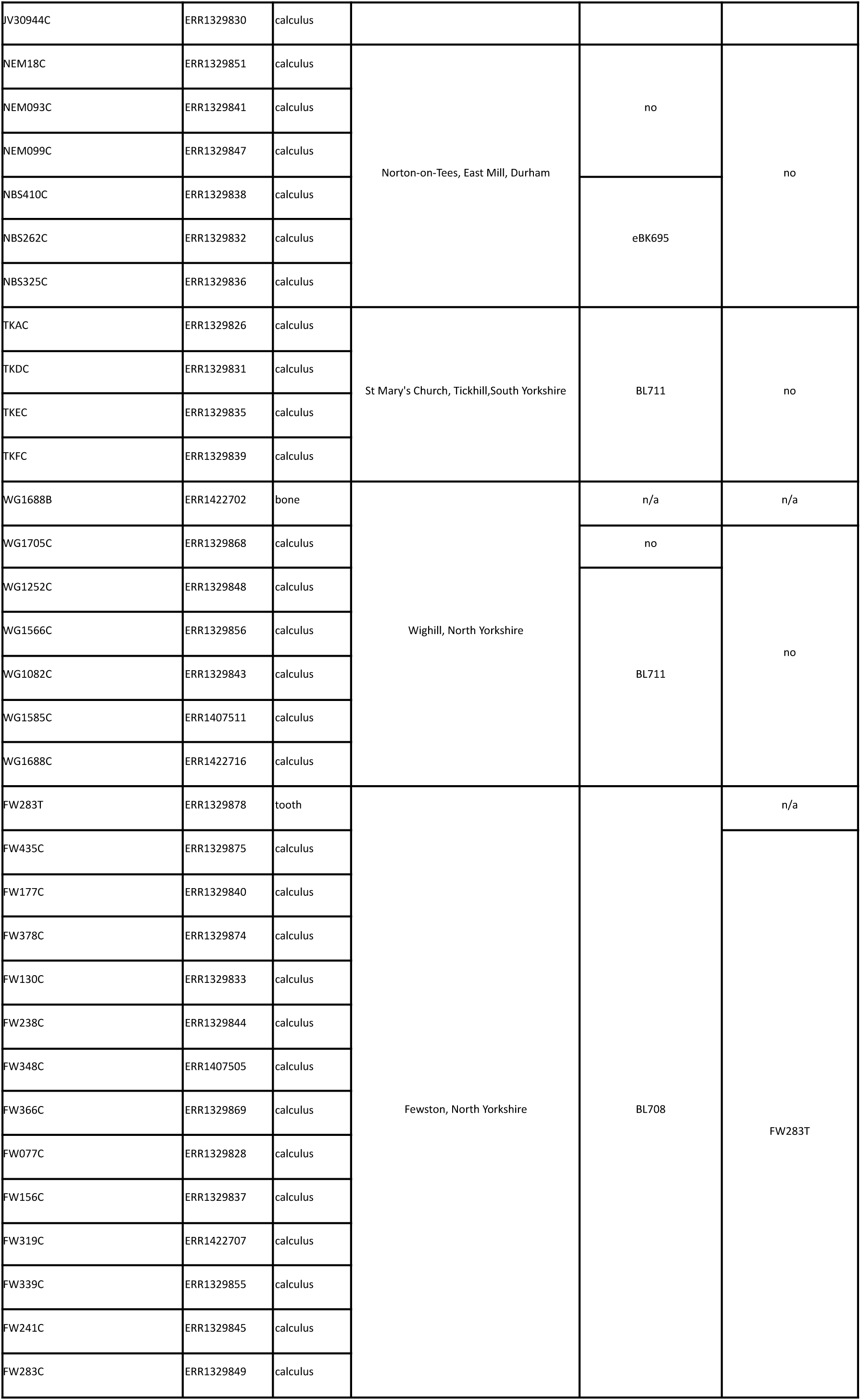
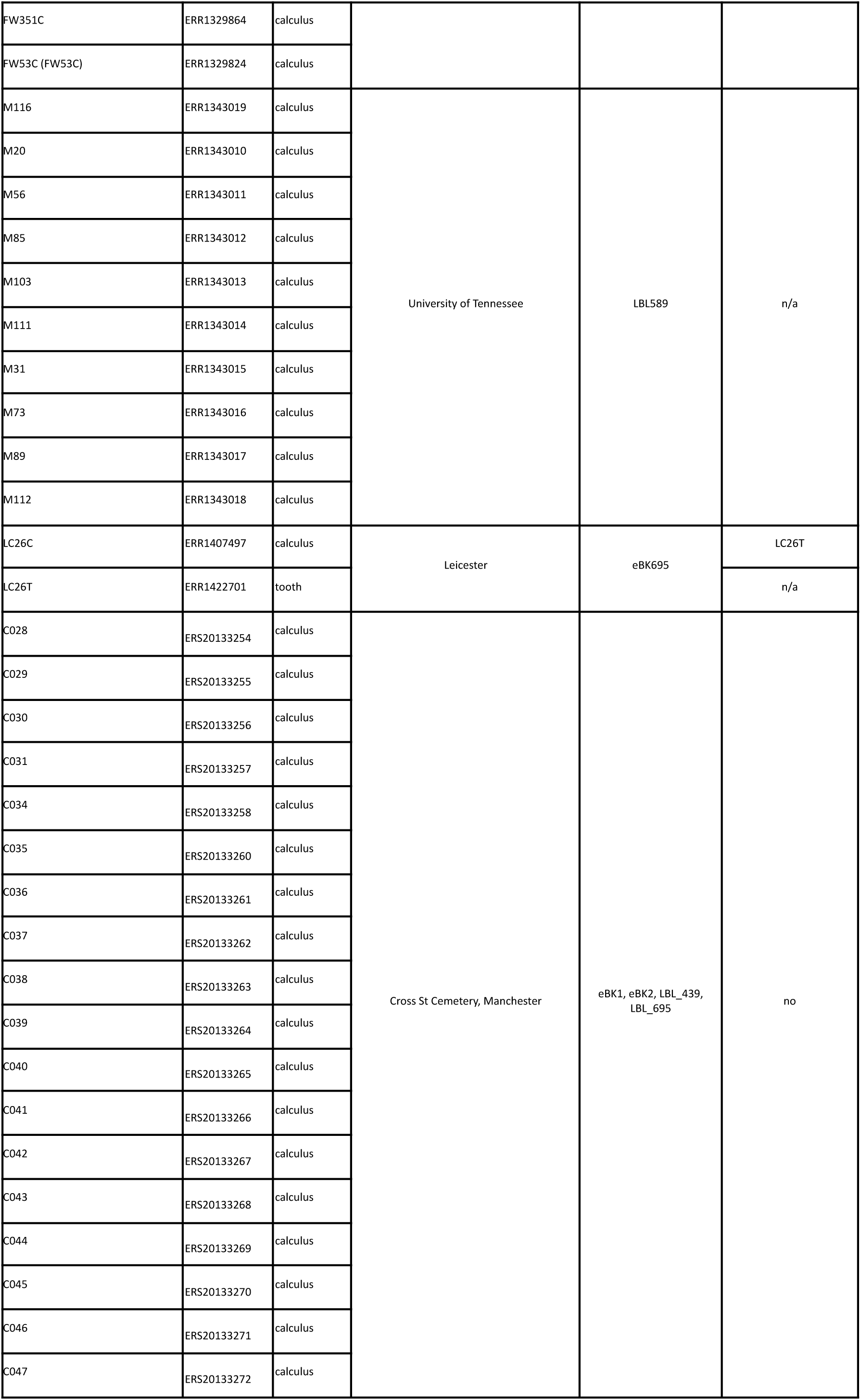
Decontamination information on samples and their matching laboratory/environmental blanks.

### aDNA authentication

Only samples (105 samples; refer to **Table 3** for this exact list) that were provided with laboratory controls (**Table 1**) were used to authenticate aDNA. The reasoning for this was to avoid potential confusion arising from modern contamination of otherwise ancient samples.

Centrifuge v1.0.3 (Kim et al. 2016) was used, with default parameters, to assign taxonomic labels by mapping sequences against the human genome, prokaryotic genomes, and viral genomes including 106 SARS-CoV-2 complete genomes. Human reads from Centrifuge outputs were retained and seqtk ‘subseq’ was used to convert them into fastq files which were then mapped to the human genome (hg38) (NCBI 2013) using BWA mem v0.7.17 (Li and Durbin 2009). SAMtools v1.12 (Danecek et al. 2021) (-view -rmdup -flagstat -sort -index) was then utilised for alignment formatting and were sorted into BAM files which were run through mapDamage2 v2.2.2 with default parameters (Jónsson et al. 2013).

## Data records

We recognize the importance of facilitating easy access to archaeological and scientific data. Analogue and digital repositories, reports and references related to skeletal individuals, and ENA projects associated with digital data in this publication can be found in **Table S1**.

## Technical validation

There are a number of concerns regarding the feasibility of our aDNA data that need to be addressed. These include absent controls, short read length, and damage authentication issues that may cause issues for potential future research projects.

### Experimental controls

It was not always possible to obtain environmental controls (such as soil, bone, or tooth root blanks) or laboratory controls (such as a library or extraction blanks) (see **Table 1**); therefore, depending on the biological question, some researchers may wish to exclude these samples from downstream analysis. It is beneficial to use environmental controls in projects that analyse the microbial composition of human microbiomes through methods such as diversity or AMR analysis. Environmental controls allow researchers to screen for and eliminate environmental contamination from their findings. These environmental operational taxonomic units (OTUs) may include the modern, living microbes at the grave site, or even the DNA from extinct microorganisms, such as ancient soil-dwelling species or those that hail from other animals and plants, that could distort the findings of a human microbiome study.

### Quality assessment

All raw and trimmed data quality, as well as read counts after trimming and decontamination can be viewed in **Table 2** and in **Figure 2**. FastQC and Fastp reports are accessible in the **supporting data (gzip folders)**.

**Figure 2.**
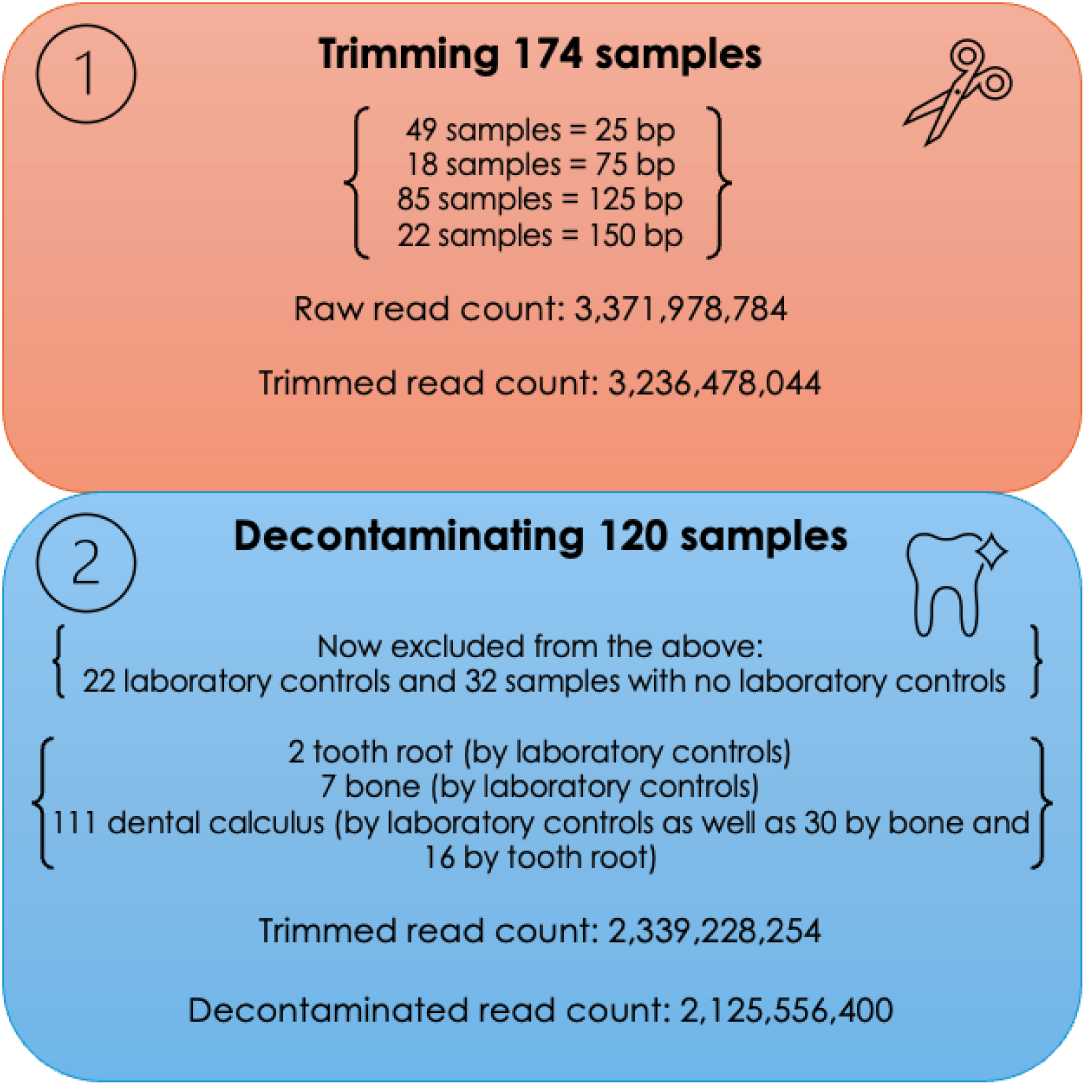
Flow chart of raw, trimmed, and decontaminated read counts.

**Table 2.** Quality control results showing percentage of mismatches sequences and reads past filters, and bp and read length pre and post-trimming and pre and post-decontamination.

| ENA/ERS<br>(specified)<br>number | Previously<br>trimmed | Mismatched<br>sequences<br>(%) | Reads<br>passed<br>filters (%) | Mean bp before<br>filtering | Mean bp after<br>filtering | Read length<br>before<br>trimming | Read length after<br>trimming | Read length after<br>decontamination |
| --- | --- | --- | --- | --- | --- | --- | --- | --- |
| 1329824 | Y | N | 99.97 | 25 | 25 | 2583158 | 2582416 | 2487082 |
| 1329825 | Y | N | 99.96 | 25 | 25 | 879712 | 879380 | 876832 |
| 1329826 | Y | N | 99.97 | 25 | 25 | 409780 | 409684 | 409684 |
| 1329827 | Y | N | 99.96 | 25 | 25 | 644972 | 644748 | 639120 |
| 1329828 | Y | N | 99.96 | 25 | 25 | 461822 | 461682 | 447704 |
| 1329829 | Y | N | 99.93 | 25 | 25 | 605826 | 605460 | 604424 |
| 1329830 | Y | N | 99.96 | 25 | 25 | 1369538 | 1369022 | 1348176 |
| 1329831 | Y | N | 99.97 | 25 | 25 | 476316 | 476216 | 476214 |
| 1329832 | Y | N | 99.97 | 25 | 25 | 892672 | 892432 | 882258 |
| 1329833 | Y | N | 99.97 | 25 | 25 | 548974 | 548856 | 514522 |
| 1329834 | Y | N | 99.96 | 25 | 25 | 388638 | 388502 | 387788 |
| 1329835 | Y | N | 99.98 | 25 | 25 | 239084 | 239058 | 239058 |
| 1329836 | Y | N | 99.89 | 25 | 25 | 391314 | 390916 | 386706 |
| 1329837 | Y | N | 99.97 | 25 | 25 | 210862 | 210818 | 201148 |
| 1329838 | Y | N | 99.94 | 25 | 25 | 198406 | 198304 | 196176 |
| 1329839 | Y | N | 99.96 | 25 | 25 | 247038 | 246944 | 246944 |
| 1329840 | Y | N | 99.97 | 25 | 25 | 467224 | 467108 | 451178 |
| 1329841 | Y | N | 99.93 | 25 | 25 | 74130 | 74084 | No match to blank |
| 1329842 | Y | N | 99.97 | 25 | 25 | 809948 | 809716 | 806872 |
| 1329843 | Y | N | 99.97 | 25 | 25 | 897988 | 897792 | 897790 |
| 1329844 | Y | N | 99.97 | 25 | 25 | 152184 | 152148 | 147296 |
| 1329845 | Y | N | 99.97 | 25 | 25 | 883494 | 883276 | 843806 |
| 1329846 | Y | N | 99.97 | 25 | 25 | 721958 | 721780 | 720434 |
| 1329847 | Y | N | 99.87 | 25 | 25 | 125702 | 125540 | No match to blank |
| 1329848 | Y | N | 99.96 | 25 | 25 | 402624 | 402492 | 402492 |
| 1329849 | Y | N | 99.82 | 25 | 25 | 251528 | 251084 | 243366 |
| 1329851 | Y | N | 99.97 | 25 | 25 | 491376 | 491238 | No match to blank |
| 1329853 | Y | N | 99.64 | 25 | 25 | 231618 | 230790 | 229678 |
| 1329855 | Y | N | 99.97 | 25 | 25 | 171824 | 171780 | 162654 |
| 1329856 | Y | N | 99.95 | 25 | 25 | 78242 | 78210 | 78210 |
| 1329857 | Y | N | 99.96 | 25 | 25 | 136876 | 136824 | 135554 |
| 1329861 | Y | N | 99.96 | 25 | 25 | 261800 | 261700 | 258868 |
| 1329862 | Y | N | 99.96 | 25 | 25 | 181530 | 181462 | 179992 |
| 1329864 | Y | N | 99.96 | 25 | 25 | 111654 | 111616 | 108182 |
| 1329865 | Y | N | 99.68 | 25 | 25 | 275252 | 274374 | 271342 |
| 1329866 | Y | N | 99.85 | 25 | 25 | 321792 | 321326 | 319042 |
| 1329867 | Y | N | 99.93 | 25 | 25 | 850018 | 849476 | 849112 |
| 1329868 | Y | N | 99.98 | 25 | 25 | 725652 | 725516 | No match to blank |
| 1329869 | Y | N | 99.97 | 25 | 25 | 269398 | 269336 | 256532 |
| 1329870 | Y | N | 99.9 | 25 | 25 | 416958 | 416556 | 413234 |
| 1329871 | Y | N | 99.96 | 25 | 25 | 450618 | 450452 | 445752 |
| 1329872 | Y | N | 99.96 | 25 | 25 | 287424 | 287336 | 283892 |
| 1329874 | Y | N | 99.46 | 25 | 25 | 19568 | 19464 | 19006 |
| 1329875 | Y | N | 99.97 | 25 | 25 | 194014 | 193966 | 186070 |
| 1329878 | Y | N | 99.97 | 25 | 25 | 394528 | 394416 | 392586 |
| 1329879 | Y | N | 99.98 | 25 | 25 | 52032 | 52026 | Blank sample |
| 1329880 | Y | N | 99.99 | 25 | 25 | 20898 | 20896 | Blank sample |
| 1329881 | Y | N | 100 | 25 | 25 | 2770 | 2770 | Blank sample |
| 1329882 | Y | N | 100 | 25 | 25 | 322 | 322 | Blank sample |
| 1343010 | N | 19.32 | 98.2 | 75 | 65 | 42560574 | 41797270 | 40603602 |
| 1343011 | N | 16.23 | 97.08 | 75 | 67 | 22883998 | 22216802 | 21063068 |
| 1343012 | N | 17.59 | 98.48 | 75 | 67 | 34442652 | 33920804 | 32940382 |
| 1343013 | N | 20.46 | 97.87 | 75 | 67 | 30978568 | 30320904 | 29360946 |
| 1343014 | N | 13.71 | 96.69 | 75 | 64 | 34561856 | 33419504 | 32140738 |
| 1343015 | N | 18.66 | 97.23 | 75 | 66 | 33328916 | 32407552 | 31712270 |
| 1343016 | N | 23.89 | 94.21 | 75 | 68 | 31635170 | 29804612 | 28942244 |
| 1343017 | N | 26.46 | 98.03 | 75 | 68 | 27706734 | 27162150 | 25949236 |
| 1343018 | N | 15.39 | 98.29 | 75 | 66 | 30454984 | 29936826 | 28909058 |
| 1343019 | N | 12.94 | 98.3 | 75 | 66 | 30692120 | 30172286 | 29213134 |
| 1343020 | N | 32.08 | 94.99 | 75 | 65 | 13175352 | 12516206 | Blank sample |
| 1343021 | N | 96.03 | 18.35 | 75 | 72 | 9036794 | 1658910 | Blank sample |
| 1407493 | N | 83.87 | 36.28 | 75 | 68 | 359280 | 130368 | Blank sample |
| 1407497 | N | 10.96 | 97.01 | 75 | 68 | 58423618 | 56682086 | 54393516 |
| 1407505 | N | 28.94 | 94.83 | 75 | 70 | 58589354 | 55565860 | 51893282 |
| 1407510 | N | 8.28 | 97.91 | 75 | 53 | 47001744 | 46022816 | 44928798 |
| 1407511 | N | 4.74 | 97.25 | 75 | 58 | 48838650 | 47499126 | 47498426 |
| 1407514 | N | 41.84 | 74.56 | 75 | 55 | 1540026 | 1148350 | Blank sample |
| 1422701 | N | 17.66 | 96.56 | 125 | 109 | 61646806 | 59530712 | 58995200 |
| 1422702 | N | 26.46 | 93.98 | 125 | 104 | 43051330 | 40460558 | No match to blank |
| 1422707 | N | 0.88 | 97.69 | 125 | 70 | 50642508 | 49474228 | 46352006 |
| 1422716 | N | 10.8 | 95.52 | 125 | 102 | 52182752 | 49847030 | 49846178 |
| 1659110 | N | 1.46 | 98.23 | 125 | 83 | 29512290 | 28992366 | No match to blank |
| 1659111 | N | 1.01 | 98.47 | 125 | 79 | 26875932 | 26467410 | No match to blank |
| 1659112 | N | 4.8 | 98.49 | 125 | 91 | 26969988 | 26565434 | No match to blank |
| 1659113 | N | 2.5 | 97.83 | 125 | 85 | 29055384 | 28427660 | No match to blank |
| 1659114 | N | 15.01 | 85.62 | 125 | 86 | 24805462 | 21238676 | No match to blank |
| 1659115 | N | 2.05 | 98.28 | 125 | 85 | 25821896 | 25378038 | No match to blank |
| 1659116 | N | 0.8 | 98.41 | 125 | 76 | 47146778 | 46400438 | No match to blank |
| 1659117 | N | 1.98 | 98.38 | 125 | 76 | 28774712 | 28310064 | No match to blank |
| 1659118 | N | 4.63 | 98.31 | 125 | 86 | 32731048 | 32180872 | No match to blank |
| 1659119 | N | 1.38 | 98.27 | 125 | 78 | 24592154 | 24166868 | No match to blank |
| 1659120 | N | 2.66 | 98.26 | 125 | 82 | 35486302 | 34872316 | No match to blank |
| 1681523 | N | 3.26 | 96.22 | 125 | 72 | 49411398 | 47545688 | 47156130 |
| 1681525 | N | 1.26 | 97.75 | 125 | 70 | 36920280 | 36093176 | 34991188 |
| 1681526 | N | 1.3 | 97.38 | 125 | 76 | 57625826 | 56120936 | 53873302 |
| 1681527 | N | 3.26 | 96.04 | 125 | 84 | 46975236 | 45116958 | 41743542 |
| 1681530 | N | 1.63 | 97.38 | 125 | 78 | 65612894 | 63896898 | 62081334 |
| 1681531 | N | 2.05 | 96.59 | 125 | 65 | 42871320 | 41409526 | 39056832 |
| 1681532 | N | 1.42 | 98.16 | 125 | 63 | 47032120 | 46171356 | No match to blank |
| 1681533 | N | 1.11 | 98.17 | 125 | 71 | 35660320 | 35009952 | No match to blank |
| 1681534 | N | 3.51 | 98.2 | 125 | 82 | 33354436 | 32756626 | No match to blank |
| 1681535 | N | 2.69 | 97.55 | 125 | 81 | 24320390 | 23726436 | No match to blank |
| 1681538 | N | 3.44 | 97.98 | 125 | 82 | 30385320 | 29773404 | No match to blank |
| 1681541 | N | 4.35 | 97.99 | 125 | 85 | 37920560 | 37159360 | No match to blank |
| 1681542 | N | 2.39 | 98.13 | 125 | 83 | 24833088 | 24370008 | No match to blank |
| 9638253 | N | 3.2 | 99.01 | 125 | 78 | 24674054 | 24431924 | 21710018 |
| 9638254 | N | 12.35 | 98.89 | 125 | 102 | 28260842 | 27947766 | 19212448 |
| 9638255 | N | 7.18 | 98.81 | 125 | 90 | 27501276 | 27174018 | 21299116 |
| 9638256 | N | 8.77 | 98.75 | 125 | 95 | 25797650 | 25487318 | 16667600 |
| 9638257 | N | 5.7 | 98.71 | 125 | 82 | 29957450 | 29572208 | 27105134 |
| 9638258 | N | 4.51 | 98.68 | 125 | 84 | 24253730 | 23935210 | 21708146 |
| 9638259 | N | 7.26 | 98.14 | 125 | 82 | 24643382 | 24185674 | 23720532 |
| 9638260 | N | 90.18 | 53.44 | 125 | 117 | 724146 | 387022 | Blank sample |
| 9638261 | N | 6.51 | 98.59 | 125 | 91 | 27624128 | 27235894 | 20130754 |
| 9638262 | N | 14.97 | 86.89 | 125 | 77 | 26588560 | 23103962 | 22367356 |
| 9638263 | N | 7.45 | 98.32 | 125 | 93 | 27257322 | 26799464 | 18403542 |
| 9638264 | N | 15.13 | 92.23 | 125 | 79 | 4642754 | 4282398 | Blank sample |
| 9638265 | N | 7.77 | 98.52 | 125 | 89 | 27644478 | 27236206 | 25596208 |
| 9638266 | N | 2.11 | 98.28 | 125 | 76 | 25996920 | 25550050 | 23611344 |
| 9638267 | N | 8.94 | 98.05 | 125 | 83 | 25031646 | 24543538 | 21289618 |
| 9638268 | N | 2.09 | 98.21 | 125 | 78 | 25575492 | 25117978 | 22180726 |
| 9638269 | N | 1.88 | 98.22 | 125 | 79 | 28529412 | 28024144 | 24710314 |
| 9638270 | N | 8.27 | 98.14 | 125 | 85 | 30716356 | 30146808 | 26628622 |
| 9638271 | N | 33.67 | 73.3 | 125 | 87 | 15952534 | 11694636 | Blank sample |
| 9638272 | N | 92.6 | 17.65 | 125 | 108 | 8723184 | 1539914 | Blank sample |
| 9638274 | N | 1 | 98.48 | 125 | 71 | 27892400 | 27470640 | No match to blank |
| 9638275 | N | 5.49 | 98.59 | 125 | 87 | 25283884 | 24928344 | No match to blank |
| 9638276 | N | 4.49 | 98.3 | 125 | 84 | 30013726 | 29503986 | No match to blank |
| 9638277 | N | 4.93 | 97.88 | 125 | 81 | 26023962 | 25473438 | 22699452 |
| 9638278 | N | 0.61 | 98.97 | 125 | 70 | 26956626 | 26681156 | No match to blank |
| 9638279 | N | 0.55 | 99.06 | 125 | 57 | 25132870 | 24896936 | 22752928 |
| 9638280 | N | 1.99 | 98.49 | 125 | 77 | 25666712 | 25280914 | 22733888 |
| 9638281 | N | 4.28 | 98.66 | 125 | 84 | 28059552 | 27684056 | No match to blank |
| 9638282 | N | 21.46 | 85.44 | 125 | 90 | 26409456 | 22564398 | 20161446 |
| 9638283 | N | 2.98 | 97.69 | 125 | 77 | 29453596 | 28774998 | No match to blank |
| 9638284 | N | 9.02 | 92.18 | 125 | 80 | 18608440 | 17154190 | Blank sample |
| 9638285 | N | 3.6 | 98.92 | 125 | 81 | 27242600 | 26924196 | No match to blank |
| 9638286 | N | 10.83 | 89.97 | 125 | 84 | 22855790 | 20564950 | No match to blank |
| 9638287 | N | 6.21 | 98.09 | 125 | 85 | 26873758 | 26362722 | No match to blank |
| 9638288 | N | 77 | 46.89 | 125 | 104 | 924370 | 433452 | Blank sample |
| 9638289 | N | 2.32 | 96.19 | 125 | 67 | 44031332 | 42353852 | 35461384 |
| 9638290 | N | 2.57 | 96.72 | 125 | 56 | 32380974 | 31321288 | 24602766 |
| 9638291 | N | 3.39 | 96.66 | 125 | 74 | 33505890 | 32388386 | 28026322 |
| 9638292 | N | 3.4 | 95.72 | 125 | 80 | 39558620 | 37867974 | 29699936 |
| 9638293 | N | 1.9 | 97.03 | 125 | 63 | 35925288 | 34858864 | 22648130 |
| 9638294 | N | 73.42 | 32.86 | 125 | 95 | 2888654 | 949470 | Blank sample |
| 9638295 | N | 87.54 | 377.77 | 125 | 106 | 1222796 | 377772 | Blank sample |
| 9638296 | N | 87.54 | 30.89 | 125 | 106 | 41936516 | 40682502 | 33970538 |
| 9638297 | N | 2.02 | 97 | 125 | 77 | 34318286 | 33304306 | 28638038 |
| 9638298 | N | 3.93 | 97.04 | 125 | 77 | 34510834 | 33516594 | 25277366 |
| 9638300 | N | 3.64 | 94.59 | 125 | 66 | 33321126 | 31521220 | 30483986 |
| 9638301 | N | 90.36 | 53.6 | 125 | 115 | 1428434 | 765698 | Blank sample |
| 9638302 | N | 13.32 | 96.2 | 125 | 104 | 33809420 | 32526006 | 32438428 |
| 9638303 | N | 2.15 | 96.39 | 125 | 65 | 34867684 | 33609404 | 31310986 |
| 9638304 | N | 5.01 | 96.89 | 125 | 85 | 38289660 | 37099046 | 33449992 |
| 9638305 | N | 2.18 | 96.82 | 125 | 77 | 35541750 | 34414316 | 32141250 |
| 9638306 | N | 12.14 | 91.74 | 125 | 81 | 3605416 | 3307704 | Blank sample |
| 9638307 | N | 3 | 97.04 | 125 | 77 | 36222640 | 35152254 | 32784300 |
| 9638308 | N | 1.62 | 97.81 | 125 | 76 | 37429770 | 36613212 | 35064998 |
| 9638309 | N | 3.33 | 95.3 | 125 | 69 | 36701602 | 34979784 | 28690092 |
| 9638310 | N | 1.9 | 96.85 | 125 | 65 | 34990762 | 33889492 | 26642234 |
| 9638312 | N | 1.86 | 97.25 | 125 | 52 | 37248422 | 36226502 | 36085818 |
| ERS20133254 | N | 1.75 | 13.45 | 151 | 78 | 13540920 | 11801754 | 10129902 |
| ERS20133255 | N | 4.11 | 9.59 | 151 | 113 | 9650924 | 9160164 | 8583678 |
| ERS20133256 | N | 13.05 | 7.23 | 151 | 92 | 7279134 | 6762296 | 4985224 |
| ERS20133257 | N | 1.13 | 10.5 | 151 | 72 | 10566408 | 9186790 | 7749908 |
| ERS20133258 | N | 2.58 | 14.29 | 151 | 96 | 14393380 | 13572660 | 12758236 |
| ERS20133260 | N | 1.65 | 12.01 | 151 | 76 | 12086216 | 10929198 | 9706028 |
| ERS20133261 | N | 2.95 | 11.03 | 151 | 89 | 11101120 | 10242942 | 9237776 |
| ERS20133262 | N | 1.76 | 11.92 | 151 | 86 | 11993372 | 10904066 | 9815586 |
| ERS20133263 | N | 1.92 | 13.11 | 151 | 82 | 13209018 | 11221758 | 9713430 |
| ERS20133264 | N | 2.38 | 11.64 | 151 | 88 | 11712516 | 10121162 | 8946904 |
| ERS20133265 | N | 6.78 | 11.33 | 151 | 103 | 11412902 | 9722646 | 8385754 |
| ERS20133266 | N | 6.78 | 9.12 | 151 | 104 | 9186792 | 8712604 | 7620040 |
| ERS20133267 | N | 7.79 | 17.64 | 151 | 102 | 17916774 | 16828530 | 15356984 |
| ERS20133268 | N | 2.42 | 12.66 | 151 | 83 | 12760300 | 11460324 | 10016358 |
| ERS20133269 | N | 3.23 | 8.64 | 151 | 72 | 8724436 | 7620118 | 6275740 |
| ERS20133270 | N | 2.06 | 12 | 151 | 94 | 12083728 | 11220760 | 10547040 |
| ERS20133271 | N | 3.14 | 94 | 151 | 20.15 | 20271626 | 18933902 | 17435362 |
| ERS20133272 | N | 7.82 | 14.55 | 151 | 116 | 14657670 | 14051062 | 13095506 |
| ERS20133273 | N | 85.45 | 98.97 | 151 | 140 | 2072654 | 2013310 | Blank sample |
| ERS20133275 | N | 33.43 | 99.13 | 151 | 101 | 1259734 | 985856 | Blank sample |
| ERS20133276 | N | 99.93 | 98.67 | 151 | 150 | 4493018 | 4433302 | Blank sample |
| ERS20133277 | N | 99.87 | 98.32 | 151 | 150 | 667232 | 655950 | Blank sample |

#### Reads

A subset of 49 samples showed high-quality base calls that had previously been trimmed prior to this research and were 25 base pairs (bp) long. Of the remaining samples, 18 samples were 75 bp, 85 were 125 bp, and 22 were 150 bp (**Table 2**). Short reads are characteristic of aDNA; however, those samples that are 25 bp are not likely to be as practical for certain research such as damage analysis (see **damage analysis** section below), screening for functional genes, contig assembly and metagenomic assembled genome creation, since they may not provide sufficient quality DNA for these type of analyses. Apart from those 25 bp samples that had already been assessed for quality, the remaining reads were trimmed and filtered for post-processing and downstream analysis. These can be viewed in the ‘per base sequence’ graphs where the X-axis shows the individual bases for reads that have been called, and the Y-axis shows the distribution values (see Fastqc reports in **supporting data**). Samples with good quality scores exhibit a blue line that continually remains high and above a distribution value of 20, which is considered a good bp call according to fastqc reports. **Figure 4** shows overall good mean sequence quality scores (see full multiQC interactive chart in **supporting data** - **compressed zip folder 3** containing **FastQC reports for trimmed data**).

The dataset initially contained a total of 3,371,978,784 raw reads which were reduced to 3,236,478,044 reads post-filtering. An average of 6% reads per sample (median of 1.84% and interquartile range [IQR] of 4.05%) were lost in the filtering process (see **Table 2** for reads lost per sample). Within the previously trimmed 49 samples, 100% of reads passed the filtering process in two library blanks (BL10 and BL11) and the rest passed >99% of reads. With the exception of laboratory controls, the remaining samples that were not previously trimmed passed >90% of reads.

32 samples could not be decontaminated as they were not assigned a laboratory blank. Samples (*n = 89*) that did not have an environmental blank were still subject to decontamination protocols using laboratory controls, but future research should take caution using these samples for further analysis as ancient and modern environmental contaminants may be present. Alternatively, using a tool like Sourcetracker (Knights et al. 2011) may help understand environmental composition in these samples to an extent. 120 remaining samples (2 tooth root, 7 bone, and 111 dental calculus) comprising 2,339,228,254 reads were all decontaminated with laboratory controls and 30 of these calculus samples were decontaminated with bone and 16 with tooth root (**Table 1**). Following decontamination, 2,125,556,400 (an average of 7.3%, median of 4.47%, and IQR of 9.99%) reads were removed (**Table 2**; **Figure 3**).

**Figure 3.**
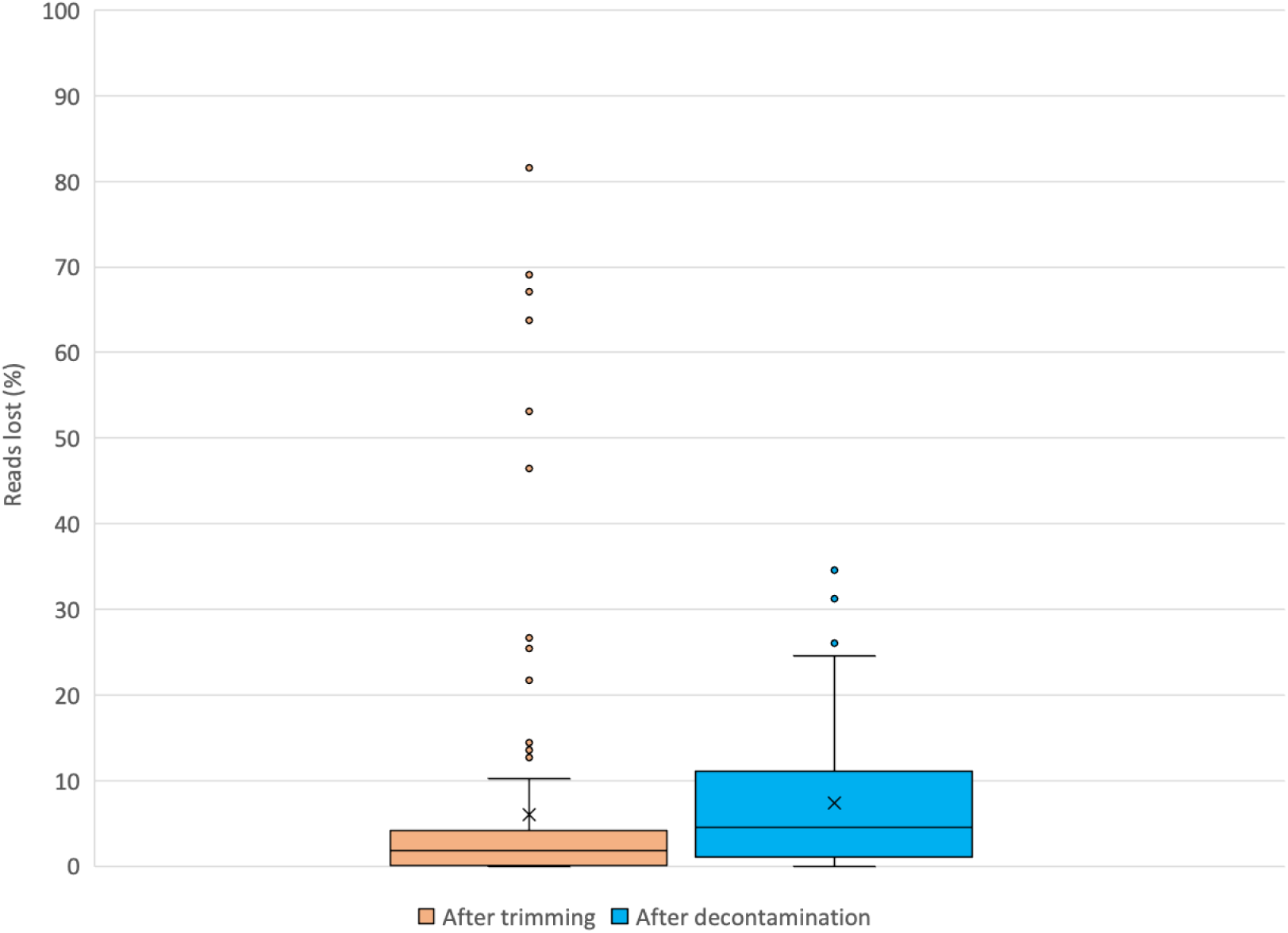
Box and whisker plots showing proportion of reads lost following trimming and decontamination.

**Figure 4.**
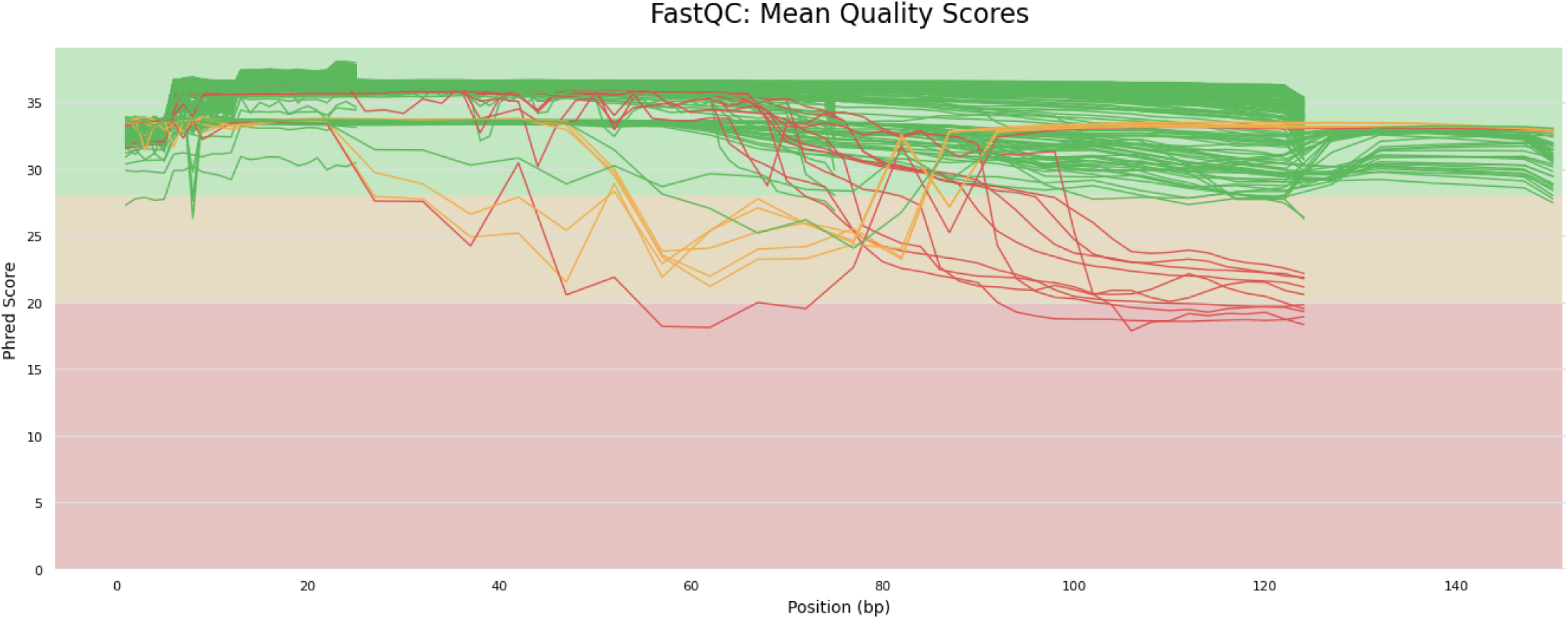
FastQC sequence quality histograms showing the mean quality value across each base position in the read for all trimmed samples. The green line represents high-quality scores (≥Q25) across all bases, the orange line indicates moderate quality scores (≥Q20-Q25), suggesting some uncertainty in the base calls, and the red line signifies low-quality scores (<Q20).

### Damage analysis

mapDamage plots (all 104 mapDamage plots are accessible in the **supplementary data** and a summary of C-T frequency transition data is shown in **Table 3**) showed varying base frequencies of C-T transitions on the 5’ ends of fragments: 36 samples produced empty plots; 59 samples show misincorporation curves below 0.025; eight showed above 0.025 but below 0.05; and one showed more than 0.05 but less than 0.10, which showed the highest levels of deamination in our sample set (**Figure 5**).

**Table 3.** Cytosine-Thymine frequency transition values for damage authentication interpretation.

| ENA/sample ID | Sample type | Average bp length | Frequency |
| --- | --- | --- | --- |
| ERR1329825 | Calculus | 25bp | empty |
| ERR1329827 | Calculus | 25bp | empty |
| ERR1329829 | Calculus | 25bp | empty |
| ERR1329830 | Calculus | 25bp | empty |
| ERR1329832 | Calculus | 25bp | empty |
| ERR1329834 | Calculus | 25bp | empty |
| ERR1329836 | Calculus | 25bp | empty |
| ERR1329838 | Calculus | 25bp | empty |
| ERR1329842 | Calculus | 25bp | empty |
| ERR1329846 | Calculus | 25bp | empty |
| ERR1329853 | Calculus | 25bp | empty |
| ERR1329857 | Calculus | 25bp | empty |
| ERR1329861 | Calculus | 25bp | empty |
| ERR1329862 | Calculus | 25bp | empty |
| ERR1329865 | Calculus | 25bp | empty |
| ERR1329866 | Calculus | 25bp | empty |
| ERR1329867 | Bone | 25bp | empty |
| ERR1329870 | Calculus | 25bp | empty |
| ERR1329871 | Calculus | 25bp | empty |
| ERR1329872 | Calculus | 25bp | empty |
| ERR1407510 | Bone | 75bp | >0.05 <0.10 |
| ERR1329826 | Calculus | 25bp | Empty |
| ERR1329831 | Calculus | 25bp | Empty |
| ERR1329835 | Calculus | 25bp | Empty |
| ERR1329839 | Calculus | 25bp | Empty |
| ERR1329843 | Calculus | 25bp | Empty |
| ERR1329848 | Calculus | 25bp | Empty |
| ERR1329856 | Calculus | 25bp | Empty |
| ERR1407497 | Calculus | 75bp | <0.025 >0 |
| ERR1407511 | Calculus | 75bp | <0.026 >0 |
| ERR1422701 | Tooth | 125bp | >0.025 <0.05 |
| ERR1422716 | Calculus | 125bp | <0.026 >0 |
| ERR1329833 | Calculus | 25bp | Empty |
| ERR1329844 | Calculus | 25bp | Empty |
| ERR1329849 | Calculus | 25bp | Empty |
| ERR1329864 | Calculus | 25bp | Empty |
| ERR1329874 | Calculus | 25bp | Empty |
| ERR9638253 | Calculus | 125bp | <0.025 >0 |
| ERR9638254 | Calculus | 125bp | <0.025 >0 |
| ERR9638255 | Calculus | 125bp | <0.025 >0 |
| ERR9638256 | Calculus | 125bp | <0.025 >0 |
| ERR9638257 | Calculus | 125bp | <0.025 >0 |
| ERR9638258 | Calculus | 125bp | >0.025 <0.05 |
| ERR9638259 | Bone | 125bp | <0.025 >0 |
| ERR9638261 | Calculus | 125bp | >0.025 <0.05 |
| ERR9638262 | Bone | 125bp | <0.025 >0 |
| ERR9638263 | Calculus | 125bp | <0.025 >0 |
| ERR9638265 | Calculus | 125bp | <0.025 >0 |
| ERR9638266 | Calculus | 125bp | <0.025 >0 |
| ERR9638267 | Calculus | 125bp | <0.025 >0 |
| ERR9638268 | Calculus | 125bp | <0.025 >0 |
| ERR9638269 | Calculus | 125bp | <0.025 >0 |
| ERR9638270 | Calculus | 125bp | <0.025 >0 |
| ERR9638277 | Calculus | 125bp | <0.025 >0 |
| ERR9638279 | Calculus | 125bp | <0.025 >0 |
| ERR9638282 | Calculus | 125bp | <0.025 >0 |
| ERR1329828 | Calculus | 25bp | Empty |
| ERR1329840 | Calculus | 25bp | Empty |
| ERR1329845 | Calculus | 25bp | Empty |
| ERR1329875 | Calculus | 25bp | Empty |
| ERR1407505 | Calculus | 75bp | <0.025 >0 |
| ERR1681523 | Calculus | 125bp | <0.025 >0 |
| ERR1681525 | Calculus | 125bp | >0.025 <0.05 |
| ERR1681526 | Calculus | 125bp | <0.025 >0 |
| ERR1681527 | Calculus | 125bp | <0.025 >0 |
| ERR1681530 | Calculus | 125bp | <0.025 >0 |
| ERR1681531 | Calculus | 125bp | >0.025 <0.05 |
| ERR9638280 | Calculus | 125bp | <0.025 >0 |
| ERR9638289 | Calculus | 125bp | <0.025 >0 |
| ERR9638290 | Calculus | 125bp | <0.025 >0 |
| ERR9638291 | Calculus | 125bp | <0.025 >0 |
| ERR9638292 | Calculus | 125bp | <0.025 >0 |
| ERR9638293 | Calculus | 125bp | <0.025 >0 |
| ERR9638296 | Calculus | 125bp | <0.025 >0 |
| ERR9638297 | Calculus | 125bp | >0.025 <0.05 |
| ERR9638298 | Calculus | 125bp | <0.025 >0 |
| ERR9638300 | Calculus | 125bp | <0.025 >0 |
| ERR9638302 | Bone | 125bp | <0.025 >0 |
| ERR9638303 | Calculus | 125bp | >0.025 <0.05 |
| ERR9638304 | Calculus | 125bp | <0.025 >0 |
| ERR9638305 | Calculus | 125bp | <0.025 >0 |
| ERR9638307 | Calculus | 125bp | >0.025 <0.05 |
| ERR9638308 | Calculus | 125bp | <0.025 >0 |
| ERR9638309 | Calculus | 125bp | <0.025 >0 |
| ERR9638310 | Calculus | 125bp | <0.025 >0 |
| ERR9638312 | Bone | 125bp | <0.025 >0 |
| ERS20133254 | Calculus | 150bp | <0.025 >0 |
| ERS20133255 | Calculus | 150bp | <0.025 >0 |
| ERS20133256 | Calculus | 150bp | <0.025 >0 |
| ERS20133257 | Calculus | 150bp | <0.025 >0 |
| ERS20133258 | Calculus | 150bp | <0.025 >0 |
| ERS20133260 | Calculus | 150bp | <0.025 >0 |
| ERS20133261 | Calculus | 150bp | <0.025 >0 |
| ERS20133262 | Calculus | 150bp | <0.025 >0 |
| ERS20133263 | Calculus | 150bp | <0.025 >0 |
| ERS20133264 | Calculus | 150bp | <0.025 >0 |
| ERS20133265 | Calculus | 150bp | <0.025 >0 |
| ERS20133266 | Calculus | 150bp | <0.025 >0 |
| ERS20133267 | Calculus | 150bp | <0.025 >0 |
| ERS20133268 | Calculus | 150bp | <0.025 >0 |
| ERS20133269 | Calculus | 150bp | <0.025 >0 |
| ERS20133270 | Calculus | 150bp | <0.025 >0 |
| ERS20133271 | Calculus | 150bp | <0.025 >0 |
| ERS20133272 | Calculus | 150bp | <0.025 >0 |

**Figure 5.**
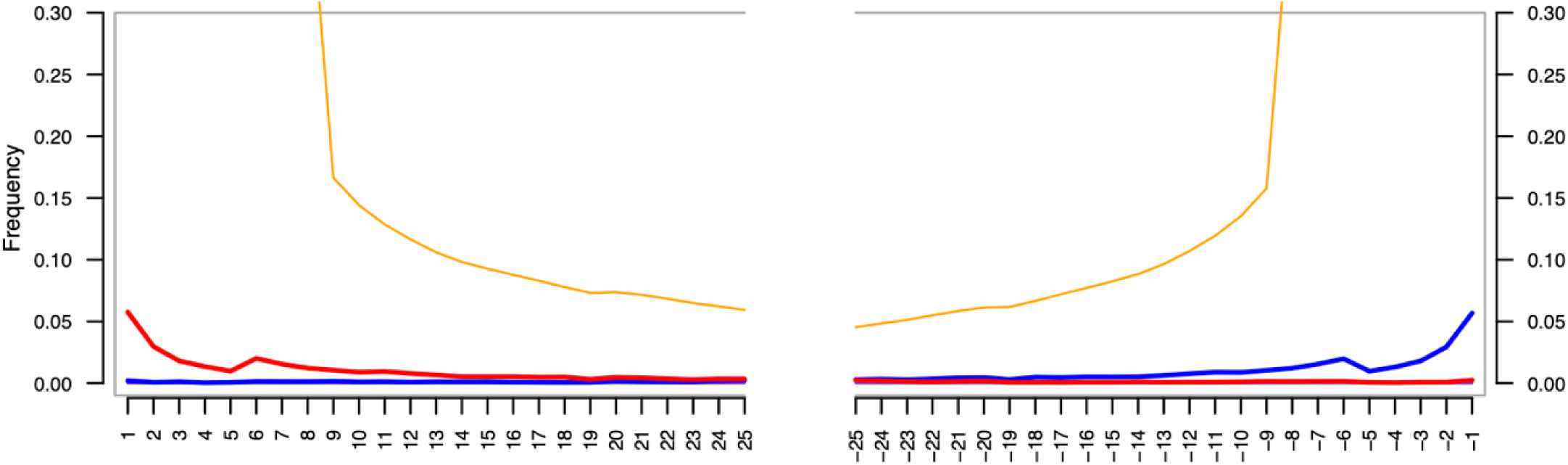
Example of a mapDamage result of C-T (5’ end) and G-A (3’ end) base substitutions from bone sample OX04B. The orange line shows nucleotides that do not align to the query sequence.

The 36 samples that returned empty plots can all be attributed to short read length (25bp). Current research does not provide a minimum nucleotide substitution frequency threshold to identify DNA as ancient, but it is clear that many of our samples (n=59) show low levels of deamination (<0.025) in contrast to other studies with C-T transition frequency curves as high as 0.3 (Andrades Valtueña et al. 2017).

There are many reasons for the low deamination levels in our ancient samples. In addition to having human reads pre-filtered by the WTSI prior to deposition in the ENA, the majority of samples contained extremely low frequencies of human genomic data because of the nature of metagenomic samples; thus, few human sequences were available to map to the human genome. Second, when mapping short, fragmented and taxonomically diverse sequences to a large and complete reference, like the human genome, mapping tools like BWA are likely to produce low coverage alignments. This observation is evidenced in **Figure 5** by the notable orange line, which would not be visible were it a high coverage alignment. These low coverage alignments make it difficult for authentication tools to identify miscoded lesions in nucleotide sequences. For example, it has been argued that reference-based mapping approaches like mapDamage (Jónsson et al. 2013) and DamageProfiler (Neukamm et al. 2021), and similar limitations in PyDamage (Borry et al. 2021), are impractical approaches for metagenomic samples when handling thousands of unidentified species (Everett and Cribdon 2023).

These tools may be more successful when aligning sequences belonging to bone or other parts of the tooth, such as the pulp, to the human genome. Our strongest damage pattern came from one of our bone samples (**Figure 5**), and other studies have retrieved high damage patterns from bone and teeth using mapDamage (Andrades Valtueña et al. 2017), whereas research that attempts to authenticate shotgun sequences from dental calculus has received results similar to ours (Gancz and Weyrich 2023). We most likely observe this pattern because dental calculus is understood to contain less endogenous human aDNA than other skeletal tissues (Ziesemer et al. 2019). Authenticating aDNA in dental calculus is still possible and has proven to be effective, for example, a 16S rRNA study by (Ziesemer et al. 2015) produced mapDamage results showing typical damage patterns in ancient dental calculus sequences. This study, however, had the advantage of using a targeted gene approach where the complexity of microbial diversity would not have impacted the reference gene alignment.

In summary, the disadvantage of applying ancient shotgun metagenomics to dental calculus, at least here, is that there are insufficient reads to produce damage signals. Although we understand that damage authentication of aDNA is essential, we think that focusing on minimising the impact of modern contamination is more important at this point.

## Summary

This article has described the preparation of an established dataset designed for research utilising microbial aDNA identified in ancient dental calculus, enabling the study of oral microbiomes from both burial populations and individuals, contributing valuable insights to archaeological science. It supports the analysis of microbial interactions in the oral cavity and includes metagenomic data from tooth roots and bone to serve as controls, helping to mitigate contamination risks. Future research projects, however, should take note of the challenges in the field of ancient metagenomics, such as absent controls, short read lengths, and damage authentication issues. Overall, our ancient dental calculus dataset offers the potential to deliver detailed insights into ancient human oral microbiomes. By making this data accessible, we aim to help advance the field of ancient DNA research and enable the exploration of archaeological topics such as diet, disease, and AMR.

## Code availability

All bioinformatic scripts are available on GitHub (https://github.com/DrATedder/dental_calculus_dataset/tree/main) for all pre-processing (quality control, decontamination) and post-processing (mapDamage) protocols.

## Author contributions

Conceptualization: C.S., S.F., M.C. and G.T., with input from ?. Supervision: C.S., A.T., C.J.M., Funding acquisition: C.S., A.T., C.J.M. Resources: M.H., A.M., C.S., G.T., M.C., and ??. Investigation: F.J.S, G D-A, J.H., C.S., S.F., K.M. Formal Analysis: F.J.S, G. D-A., J.H., C.S., S.M., A.T. and J.W. Data Curation, Visualization and Writing - Original Draft: F.J.S, G. D-A. Writing - Review & Editing: all authors.

## Acknowledgments

We thank the following individuals, museums and agencies for providing access to skeletal collections: Forensic Anthropology Center at the University of Tennessee, Knoxville (Dawnie Steadman); John Buglass Archaeological Services; Museum of London (Rebecca Redfern, Jelena Bekvalac); Natural History Museum (Ian Barnes, Heather Bonney); Durham University Department of Archaeology (Anwen Caffell, Rebecca Gowland); University of Leicester Archaeological Services; University of York Department of Archaeology (Cath Neal); Washburn Heritage Centre; York Archaeological Trust for Excavation and Research Ltd (Christine McDonnell); York Osteoarchaeology Ltd (Katie Keefe). We are grateful to Dr Lisa MacKenzie-Davey for helping to sample the calculus from Manchester Cross Street. Thanks to Julian Parkhill, Josef Wagner and David Jackson from the Wellcome Trust Sanger Institute for assistance with project design and metagenomic sequencing. We are grateful for the assistance of Drs Gavin Thomas and Sandy McDonald (University of York) for their advice on research design, and to Drs Mark Jenner and Sarah Goldsmith (University of York) for input on the historical context of the post-medieval skeletons. The authors acknowledge the use of the University of Bradford High Performance Computing Service in the completion of this work.

## Table legends

**Table S1.** Skeletal information, proveniences and ENA accession numbers for the archaeological and modern dental calculus samples collected in this study.

## Supporting data

Table S1 (large metadata table)

Compressed zip folder 1: FastQC reports for raw data

Compressed zip folder 2: Fastp reports

Compressed zip folder 3: FastQC reports for trimmed data

Compressed zip folder 4: mapDamage plots

